# Gene expression profiles of some prognostic markers in a human breast cell line (MCF7) exposed to Curcumin nanoparticles and nanocapsules

**DOI:** 10.1101/2022.01.04.474885

**Authors:** Emad S. Shaker, Ghada M. Nasr, Mahmoud Moawad

## Abstract

1.

**Introduction:** Worldwide, cancer is a significant public health problem. Curcumin exhibits anti-inflammatory, antiproliferative, and anticancer properties when used in medicine. Investigated study for Curcumin’s chemopreventive mechanism against human malignancies, this research examined the cellular and molecular alterations generated by curcumin modified compound in breast cancer (MCF-7) cell lines. Oncogenic EGFR and VEGFR2 mutations lead to the formation, invasion, and maintenance of malignant phenotypes in humans, including breast cancer. Studied prognostic markers such as C-myc and Ki67 in breast cancer, and the apoptotic gene as Caspase-3 have been done.

**Aim of the work:** The purpose of this study is to determine the therapeutic efficacy of curcumin nanoparticles and nanocapsules in breast cancer cell lines (MCF7).

**Materials and methods:** We used real-time PCR to assess the expression of the C-myc, Ki67, EGFR, VEGFR2, and Caspase-3 genes in MCF7 cells treated with Curcumin nanoparticles and nanocapsules.

**Results:** Curcumin nanoparticles and nanocapsules boosted apoptotic cell populations considerably regardless of the nanotechnology used. Additionally, the mRNA expression analysis results indicated that the mechanism activated by curcumin nanocapsules involved the upregulation of the oncogenes EGFR and VEGFR2. In comparison to curcumin nanoparticles, curcumin nanocapsules significantly reduced the expression of Ki67 and c-myc mRNAs in breast cancer cells. The mRNA expression study revealed that curcumin nanocapsules produce an increase in the apoptotic Caspase-3 gene production compared to cells treated with curcumin nanoparticles.

**Conclusion:** This work demonstrates that curcumin nanoparticles created using a novel mechanical process can be employed successfully as an anticancer agent. These findings add to our understanding of the molecular mechanisms behind curcumin nanocapsules’ anticancer activity in breast cancer.

## 2. Introduction

For millennia, cancer has posed a significant threat to public health, and its prevalence is increasing at an alarming rate. According to worldwide cancer statistics (GLOBOCAN) 2020, cancer incidence will grow by around 47% in 2040 compared to predicted cases in 2020. The increasing number of cancer cases will almost certainly increase the death rate of cancer patients, particularly breast and colorectal cancer [1]. Breast cancer is the second most prevalent malignancy and the sixth most lethal. Breast cancer accounts for 23% of all female cancer diagnoses and 14% of cancer deaths. Breast cancer is a term that refers to a group of cancers that arise from the breast epithelial cells. Cell lines can be utilized as *in vitro* models for cancer research in molecular diagnostics [2].

The EGF receptor (EGFR) controls cell growth, including HER-1, EGFR-2/HER-2, HER-3, and HER-4 (Erb-4). EGFR-1 and HER-2 are overexpressed in breast cancer [3]. The most common EGFR-1 ligands are EGF and TGF, which form complexes with HER-2 and other ErbB family receptors. Binding and dimerization activate the intracellular tyrosine kinase [4]. Activated EGFR (pEGFR) promotes MAPK/ERK and PI3K/Akt signaling pathways, causing uncontrolled cell proliferation and apoptosis suppression. Other prognostic factors include the activated EGFR variant (pEGFR) [5]. EGFR phosphorylation has been associated to a poor prognosis in non-small cell lung cancer [6]. Oncogenic EGFR overexpression requires medicines that disrupt EGFR signaling through phosphorylation and its target molecules.

sVEGFR-1 is a soluble receptor tyrosine kinase (RTK) that functions as a negative regulator of VEGF [7]. VEGFR-2 is a VEGF/VEGFR angiogenic signaling receptor. Breast cancer must form localized tumors and spread [8]. Microvessel counts relate higher VEGF expression to poor prognosis in several malignancies, including breast [9].

VEGF controls cancer angiogenesis and cell growth [10]. However, clinical trials have shown that additional inhibitors may be required to control tumor development and proliferation. They employ the same downstream signaling molecules. Tumors with constitutive VEGF overexpression acquire resistance to anti-EGF/EGFR treatment [11]. Activating EGFR signaling increases VEGF production. Because both routes are activing at the same time, combining EGF(R) and VEGF(R) inhibition may have anticancer effects. Combining EGFR and VEGFR inhibitors seems to slow tumor development [12]. In colorectal, breast, and lung cancer trials, bevacizumab (humanised anti-VEGF antibody) showed promise [13].

The determination of the proliferative fraction of cancer cells using Ki-67 expression was a simple, reproducible procedure. The expression of Ki-67 was examined in several different types of human cancer. Ki-67 was favorably linked with tumor grade in breast cancer [14]. MYC proteins are transcription factors that regulate gene expression and cancer. Myc family member overexpression is expected in over half of all human cancers. C-myc is the family’s defining member, overexpressed in hematologic and solid tumors [15].

Caspase 3, Bax, and Bcl2 regulate apoptotic cell death and proliferation [16]. Caspase-3 is an endoprotease that controls apoptosis and inflammation. Apoptotic executioner caspase-3 coordinates the destruction of cellular structures such as DNA fragmentation and protein cytoskeletal disintegration. [17].

Nanomedicine is a rapidly growing science that can significantly alter human health [18]. Our group studied curcumin micro and nanocapsules in generated considerable investigation as antioxidant and anticancer agents [X, 20]. Curcumin has gained increased interest for its antibacterial, antioxidant, and anti-aging properties, and its ability to scavenge free radicals. Numerous further researches have credited Curcumin’s broad spectrum of therapeutic qualities, [21], including its ability to inhibit cancer cell proliferation, promote tumor cell death, and prevent metastasis [22]. Curcumin was proved to decrease the proliferation of cancer cells in various cancer types, including gastric, hepatic, breast, colon, cervical, and lung cancer [23]. Additionally, curcumin administration significantly alleviates the adverse effects associated with traditional therapy [24].

This study was designed to determine the therapeutic effect of curcumin nanoparticles (CNP) and Curcumin nanocapsules (CNC) on a breast cancer cell line (MCF7) as well as the expression patterns of the EGFR, VEGFR2, C-myc, Ki 67, and caspase3 genes.

## 3- Material and Methods

### Preparation of curcumin nanocapsules

Curcumin nanocapsules CNC were made by mechanically milling (Model: PQ-N2 Planetary Ball Mill, Gear Drive 4-station planetary Ball mill, 220v) [25] at 40,000 rpm for 90 minutes. A magnetic stirrer was employed to extract pulverized Curcuma powder with 95% ethanol [26]. The CNC manufacture used homogenization, sodium alginate and Tween 20 (T20) matrix [19].

### Transmission Electron Microscopy TEM

CNC was analyzed using TEM to determine its morphology. After accelerating the voltage and observing the specimens under a microscope, the diameter of the specimens was estimated using micrographs [27].

### Molecular Studies

In breast cancer cell lines [MCF7], we examined the mRNA expression of the EGFR and VEGFR2 genes in combination with B-actin as a housekeeping gene (control gene) implicated in the pathway. The RNA was isolated from the cells using an RNA extraction kit (Quigene). The RNA quality was assessed using agarose gel electrophoresis. NanoDrop 1000 Spectrophotometer assessed optical density (A260/A280 ratio) of isolated RNA (Wilmington, DE, USA). In this case, the REVERTA-L RT reagents kit was employed for cDNA conversion.

The ExicyclerTM96 Bioneer equipment was used to perform real-time PCR using the SYBR Green dye (KapaBiosystems, Inc., Wilmington, MA, USA) (Bioneer Corporation, Daejoen, Korea).

One gram RNA was reverse transcribed using the qScript cDNATM SuperMix according to the manufacturer’s instructions.

The resulting cDNA was then subjected to RT-PCR using the primers specified in Table (1) as follows: 5 minutes at 95°C, 40 cycles of denaturation for 1 minute, annealing for 45 seconds at 60°C, extension for 30 seconds at 72°C, and a final extension for 20 seconds at 72°C [28, 29, 30, 31].

**Table 1:**
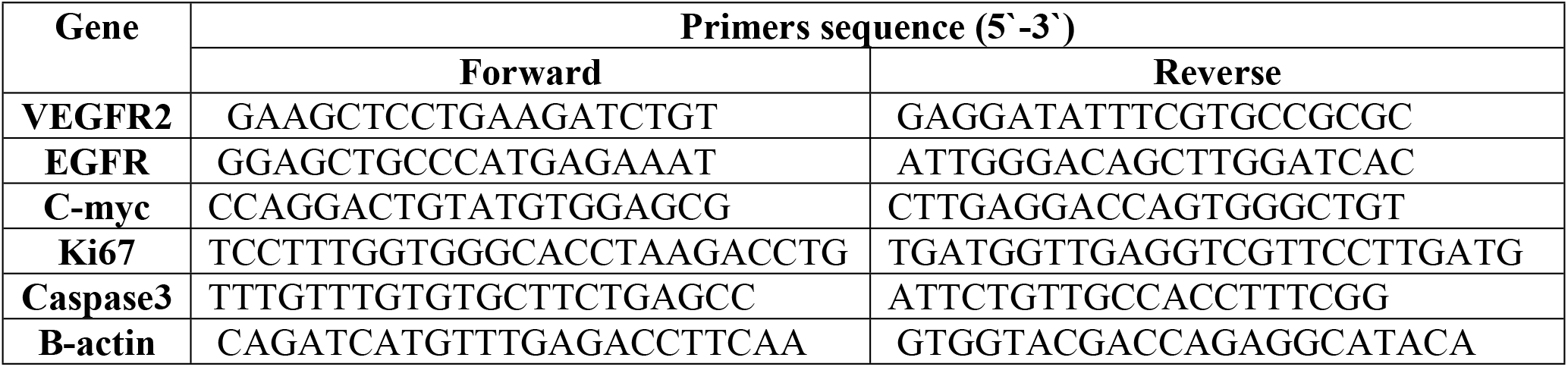
Primers for all examined genes.

### Statistical Analysis

The SPSS statistical software evaluated the data, followed by Waller-Duncan ratio [32].

## 4- RESULTS5

### Characterization of curcumin nanocapsules

TEM was used to image curcumin nanocapsules (Figure 1). The images of spherical micelles indicated that nanocapsules have a diameter ranging from 7 to 42 nm [19]. The particle size distribution of curcumin nanoencapsulation was obtained and analyzed numerically.

**Figure 1.**
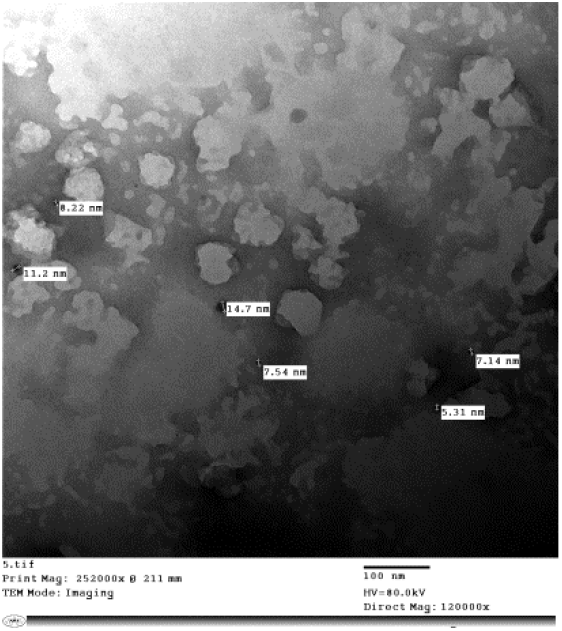
TEM images of Curcumin nanocapsules

### Quantitative Real-Time Polymerase Chain Reaction

The fold change in C-myc, Ki67, EGFR, VEGFR2, and Caspas3 mRNA expression levels in curcumin nanoparticles and nanocapsule-treated MCF7 cells compared to untreated cells is shown in Figures (2, 3, and 4). The present study demonstrated lower expression of the EGFR and VEGFR2 genes in MCF7 cells treated with curcumin nanocapsules compared to MCF7 cells treated with curcumin nanoparticles (figure 2).

**Figure 2.**
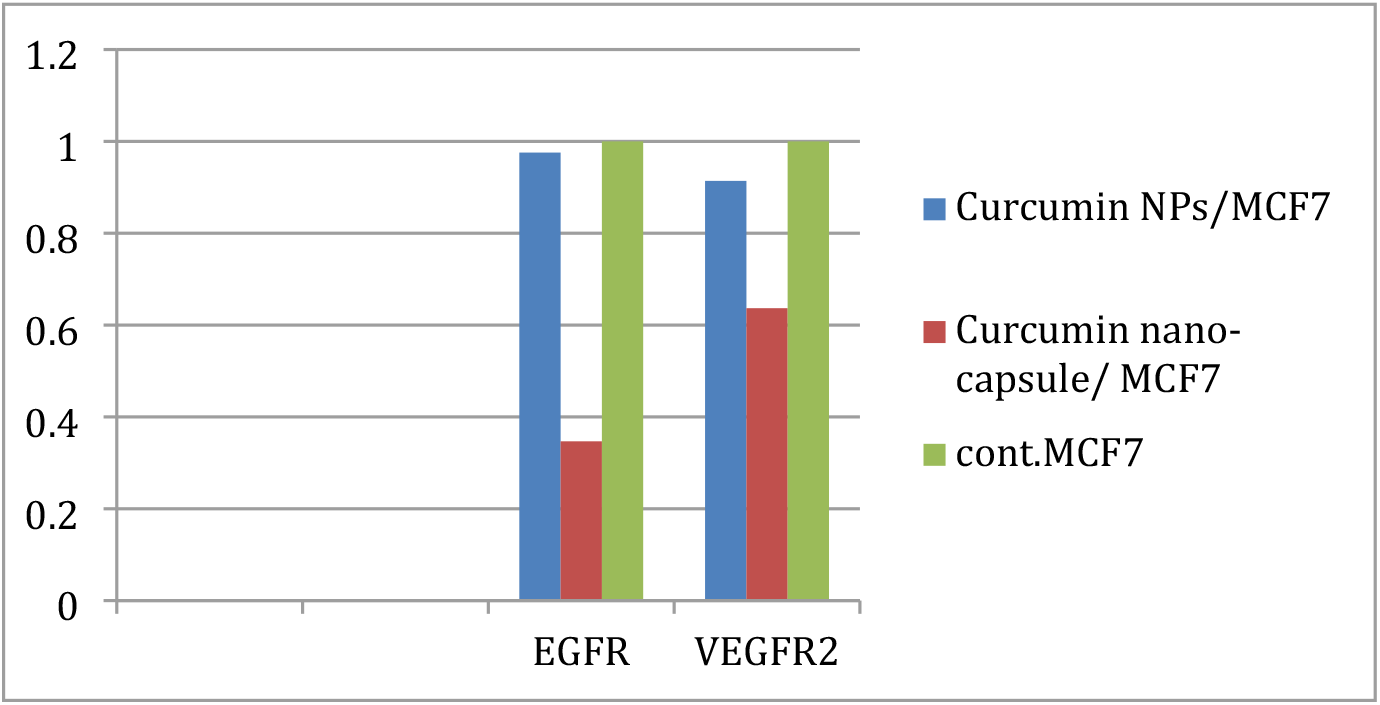
Effect of Curcumin NPs and nanocapsules on VEGFR2 and EGFR genes expression

As shown in Figures, curcumin nanoparticles treated with human breast cancer cells (MCF-7) expressed more Ki67 and C-myc mRNA than curcumin nanocapsules treated with human breast cancer cells (MCF-7) (figure 3). The fold change of the apoptotic gene caspase-3 and the housekeeping gene-actin as a control in MCF-7 cells treated with curcumin nanoparticles and curcumin nanocapsules compared to untreated cells is shown in Figure (4). When cells treated with curcumin nanoparticles or curcumin nanocapsules were compared to controls, the expression of the caspas3 gene was enhanced. On the other hand, the expression of the caspase3 gene rose considerably following treatment with curcumin nanoparticles. In control RT-PCR analysis, Curcumin did not affect the levels of β-actin mRNA in either cell line.

**Figure 3.**
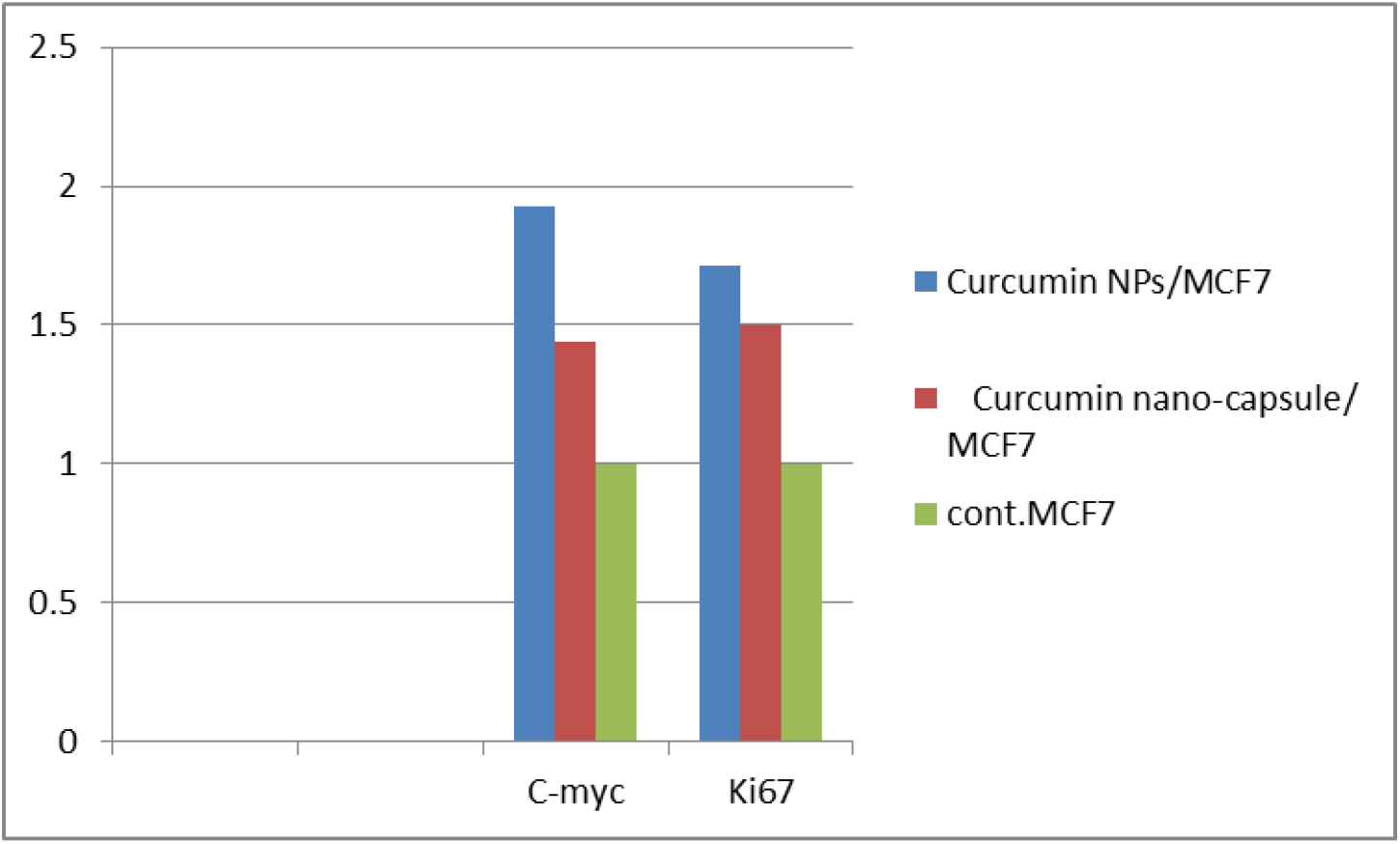
Effect of Curcumin NPs and nanocapsules on C-myc and Ki67genes expression

## 5- DISCUSSION

Locally advanced breast cancer accounts between 30% and 60% of cases with breast cancer and gradually will be a clinical challenge because most patients with this diagnosis acquire distant metastases after receiving suitable and effective radiation and surgery [33]. Locally advanced breast tumors are frequently accompanied with increased EGF and VEGF expression, decreased relapse-free or overall survival, and disease aggressiveness [34]. Thus, a non-toxic, more effective, and new treatment method is required to combat loco-regional breast cancer recurrence, particularly in individuals treated prior to chemo or radiotherapy.

EGF, VEGF, and their receptors, EGFR and VEGFR, have all been involved in the development of human cancer for an extended time [35]. These variables have solid connections and signaling, resulting in survival and resistance mechanisms that effectively evade targeted therapy [36]. EGFR is a member of the human EGF receptor (HER) family of tyrosine kinases, whose aberrant signaling has been linked in a vast variety of epithelial-derived malignancies, which account for nearly 80% of all solid tumors. [37]. EGFRs, being one of the most extensively studied receptor tyrosine kinase families, play critical roles in signal transduction and tumorigenesis. Additionally, several cancers exhibit deregulation of the EGFR gene [38]. VEGFR2 receptors were initially assumed to be exclusive to endothelial cells, but recent investigations have revealed that VEGFR2 is widely expressed in human tissues and malignancies, implying an autocrine signaling loop between VEGF and VEGFR2 [39].

Breast cancer is not the only type of cancer that is well-known. Two molecular subtypes have been identified using gene expression profiling, with diagnosis based on the presence or lack of the hormone receptor-related genes EGFR and VEGFR2. Pro-angiogenic factors are overexpressed in several types of malignancies. MCF7 breast cancer cells, one of the most aggressive kinds of cancer, have been discovered to upregulate EGFR and VEGF/VEGFR expression, both pro-angiogenic factors. The EGFR and VEGFR2 signaling pathways have been found to interact in tumor formation, survival, and angiogenesis. ERK and PI3 K/Akt signaling are activated by EGFR and VEGFR2 [40]. This regulation is shown in cancer cells that express EGFR and VEGFR2 [41]. Because we used human ramucirumab in our xenograft trial, it seems improbable that the ramucirumab arm of anti-EGFR/VEGFR2 monoclonal antibodies inhibited angiogenesis. Anti-EGFR/VEGFR2 BsAbs have anticancer efficacy by blocking EGFR signaling and altering VEGF/VEGFR2-mediated autocrine/paracrine pathways, which is consistent with our results in cellular models. To ascertain the effect of CNPs and CNCs, we analyzed the oncogene repressors EGFR and VEGFR2 to discover the mechanism by which they augment the effect of CNPs and CNC; we also conducted some tests on the expression of genes implicated in the oncogenic pathway.

In contrast to individual exposure, real-time PCR analysis demonstrated that co-treatment with CNP and CNC leads to a significant drop in EGFR and VEGFR2 expression levels in curcumin nanoparticles and nanocapsule treated cells compared to the control untreated cell line. The current study showed that when MCF7 cells were treated with CNPs, EGFR and VEGFR2 expression was increased compared to when MCF7 cells were treated with Curcumin nanocapsules. Our findings are consistent with those reported by Sarkar et al. [42].

The proto-oncogene c-myc is known to influence tumor formation and apoptotic cell death. A previous study indicated a link between p53 mutations and c-myc activation [43]. Ki67 was first identified in 1983 as a nuclear protein in a Hodgkin’s lymphoma cell line. The expression of Ki-67 was examined in some different types of human cancer. Ki-67 had a positive correlation with breast cancer grade [44].

Intracellular caspases, a class of structurally related cysteine proteases discovered previously [45], are key signaling channels involved in apoptotic cell death. Caspase activity is involved in the cleavage of cellular proteins that are often proteolyzed during apoptosis, either directly or indirectly. Caspases -2, -3, -6, -7, and -9, for example, can cleave poly (ADP ribose) polymerase (PARP) [46]. Curcumin was reported to boost the expression of downstream cleaved caspase 3, indicating that it may stimulate the mitochondrial apoptotic pathway [47].

Curcumin is a highly effective anticancer drug that acts differently on different cells. Curcumin’s capacity in causing apoptosis for various cancer cells, made scientists expect that one day curcumin derivatives be developed as universal anticancer treatment. Curcumin, a naturally occurring polyphenol chemical derived from the annual herb Curcuma longa, has been demonstrated to effectively treat malignant and benign tumors, inflammation, and a variety of other ailments [48].

This work was designed to examine the molecular mechanisms underlying the EGFR, VEGFR2, C-myc, KI67, and caspase3 gene pathways in a human breast cancer cell line (MCF7) treated with curcumin nanoparticles and nanocapsules. Scientists can create microscopic particles using nanotechnology. When nanoparticles are smaller than 20 nm or 50 nm, they can easily pass through blood vessel walls and even into most body cells, making them an attractive candidate for specialized pharmaceuticals employed in the targeted delivery of large doses of chemotherapeutic agents [49]. TEM was used to image curcumin nanocapsules. The images of spherical micelles indicated that nanocapsules have a diameter ranging from 7 to 42 nm [19, 50] and described free curcumin powder as having an irregular form and a mean particle size of around 3.58 m. Our research found that mechanical nano preparation reduced the size of curcumin powder by around 100 times, enhancing the drug’s anticipated anticancer efficacy. Curcumin nanocapsules had a smooth surface and a spherical shape, comparable to those discovered in a previous study for curcumin nanoparticles [51].

To better understand the molecular alterations generated by curcumin administration, we used RT-PCR to examine the expression of Ki67, C-myc, and caspase3 mRNA. In breast cancer cells, curcumin nanoparticles and nanocapsules treatment boosted the expression of Ki67, C-myc, and Caspas3 mRNAs, compared to untreated control cells (figures 3, and 4). In MCF-7 cells, treatment with curcumin nanocapsules decreased Ki67 and C-myc mRNA levels while increasing up-regulation compared to treatment with curcumin nanoparticles. Our findings corroborate previous findings [52, 53]. Finally, it can be stated that Curcumin nanoparticles synthesized using novel mechanical processes can be used successfully as anticancer drugs. These findings add to our understanding of the molecular mechanisms behind curcumin nanocapsules’ anticancer activity in breast cancer cells.

**Figure 4.**
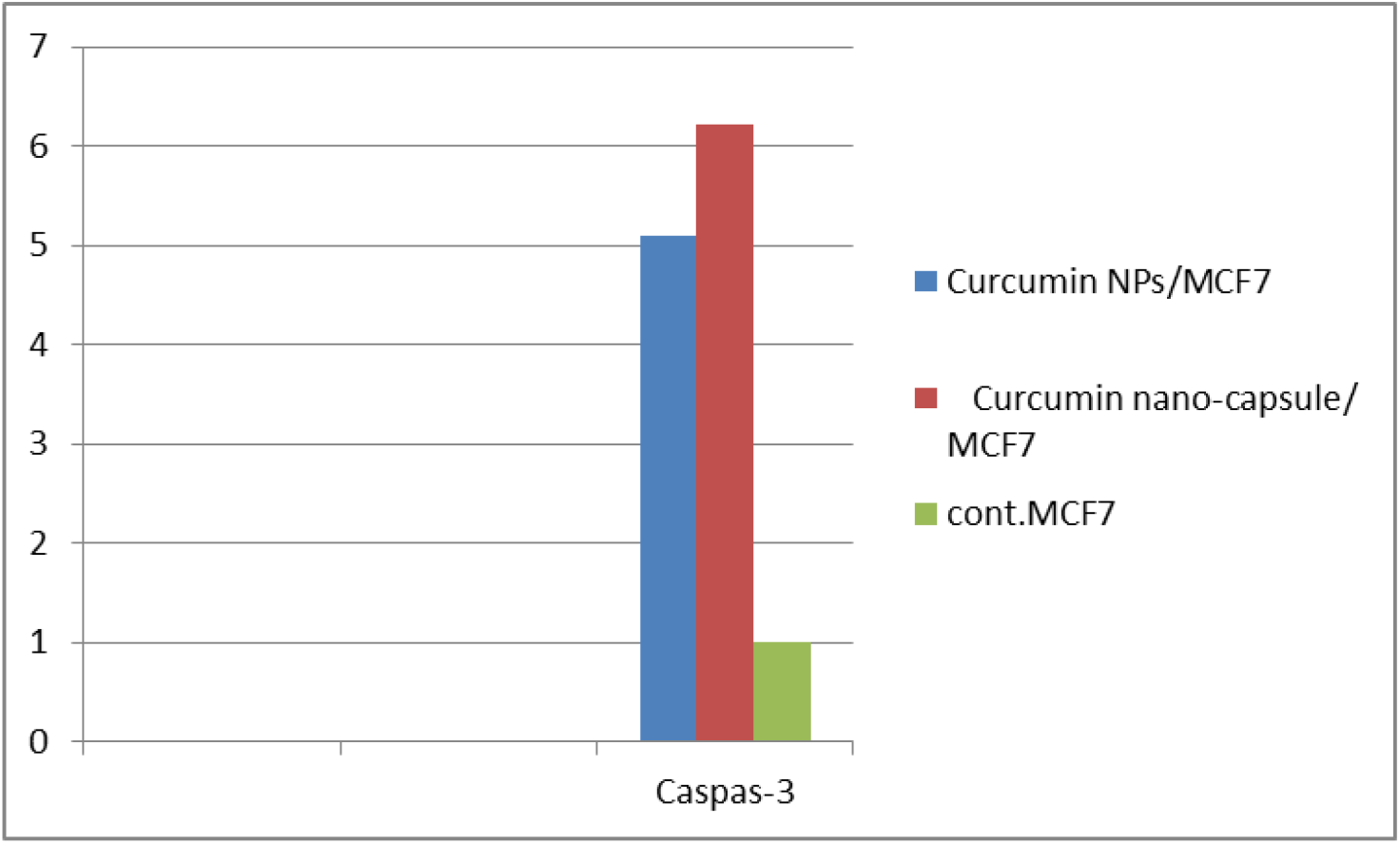
Effect of Curcumin NPs and nanocapsules on Caspase-3 gene expression

